# No effect of menstrual cycle phase on eccentric exercise-induced neuromuscular impairments and the magnitude of the repeated bout effect

**DOI:** 10.1101/2024.06.10.598231

**Authors:** Amelia Rilling, Geoffrey A. Power

## Abstract

Neuromuscular function is impaired following an unaccustomed bout of eccentric exercise, however, through the repeated bout effect (RBE) the muscle is protected from damage following a subsequent bout of eccentric exercise. As a result of unaccustomed eccentric contractions, structural muscle damage occurs in both sexes. However, the inflammatory response may be mitigated in females due to estradiol, thereby attenuating the secondary effects of muscle damage and potentially limiting the magnitude of the RBE. We investigated the relationship between menstrual cycle phase and oral contraceptive use on neuromuscular impairments following the first bout of exercise, and the magnitude of the RBE. Thirteen female participants performed two bouts of 150 maximal eccentric voluntary contractions of the elbow flexors four weeks apart. Normally menstruating females participated during the late follicular phase (day 10-14) of their menstrual cycle, as determined through cycle tracking, when estradiol is near peak, and progesterone is lower. Oral contraceptive users were tested on their placebo pill days (lower estradiol). Neuromuscular function was assessed for Bout 1 before the eccentric protocol and then again 48 hours following, and this was repeated 4 weeks later for Bout 2. Eccentric exercise-induced muscle weakness and soreness did not differ between groups following Bout 1 (p=0.885), and the magnitude of the RBE (p<0.05) was similar between groups (p=0.995). Females in the late follicular phase (classified as high estradiol) and females on combined oral contraceptives (low estradiol) had similar impairments in neuromuscular function following the first bout of eccentric exercise, and a similar RBE.

## 1. Introduction

Sex-differences in sport have become a rapidly advancing area of research. To carefully interpret differences that arise between the sexes, further research needs to be conducted on females to understand performance, strength, and recovery variability throughout the female menstrual cycle [1]. It is known that unaccustomed active lengthening muscle contractions (i.e., eccentric exercise) results in muscle damage [2]. Muscle damage can be categorized into primary and secondary damage. Primary damage results from the initial mechanical perturbation and a structural disarray of myofibrillar machinery and is comparable between sexes [2,3]. However, females often demonstrate reduced muscle damage and a faster recovery of muscle function following eccentric exercise in both animal and human models [3,4]. This blunted damage and faster recovery in females is likely owing to an altered inflammatory processes and attenuation in secondary mediators of damage, offering protection from further damage as compared with males.5

Skeletal muscle damage following a bout of eccentric exercise offers a protective effect against successive bouts of the same exercise. This phenomenon is referred to as the repeated bout effect (RBE) [6]. The magnitude of the RBE is relative to the severity of muscle damage incurred during the initial bout of exercise [7]. The RBE has been observed in both males and females, however, limited research has examined the RBE in females or the intra-variability within females due to hormone fluctuations throughout the menstrual cycle [7,8]. A study by Bruce et al. (2021) investigated the RBE in males and females following 200 maximal eccentric contractions of the dorsiflexors, and found a similar RBE for both sexes. While there was no effect of sex, MVC torque in the female group appears to have near fully recovered by 48 hours following the eccentric protocol, indicating that the decrease in torque was likely due to fatigue and not necessarily muscle damage [9]. Their study did not account for menstrual cycle phase, offering a wide variation in estradiol levels in the female group [9]. Therefore, the literature pertaining to the impact of menstrual cycle phase (e.g., estradiol) on the magnitude of the repeated bout effect remains unclear.

Estradiol is a fluctuating hormone in females which contributes to a reduction of inflammation and leukocyte infiltration, and thus the factors responsible for the secondary effects of muscle damage after an initial bout of eccentric exercise [5]. Estradiol mediates neutrophil accumulation at the site of damage, alleviating further structural damage through two proposed methods [5,10]. The first method is through the reduction of calpain [11]. Calpain mediates the structural changes in neutrophils allowing them to move between endothelial cells to the site of muscle damage [12]. Secondly, estradiol activates endothelial nitric oxide synthase (eNOS) through phosphorylation [10]. eNOS modulates the homeostasis of endothelial cells, allowing less neutrophil infiltration through endothelial cell gaps following eNOS phosphorylation [13]. Compelling evidence by MacNeil *et al*. (2009) demonstrated a reduction in neutrophils 48 hours after eccentric exercise-induced muscle damage in males taking estradiol supplementation. Males were supplemented with estradiol for eight days to obtain a serum level of approximately 94 pg/ml (vs. 38 pg/ml for controls) [10]. Despite dramatic differences in neutrophil attenuation between groups, the levels of estradiol in these males remained substantially lower than levels found in females, thus offering potential for a more profound effect within females. Additionally, levels of caveolin-1, a lipid binding protein, were diminished after 48 hours in the experimental group [10]. Elevated levels of caveolin-1 post exercise inhibits repair processes [14]. Therefore, it is likely that estradiol alleviates further inflammation and the secondary effects of damage through neutrophil mitigation.

Additionally, inflammatory markers of muscle damage and reductions in knee extension torque following 240 maximal eccentric knee extensions were measured in males, normally menstruating females, and females on oral contraceptives [15]. The oral contraceptive group was determined to have lower overall estradiol levels than the normally menstruating females [15]. Following the eccentric protocol, knee-extension torque was reduced similarly in all groups. However, torque continued to decline 48 hours after the eccentric protocol in both the males and females on oral contraceptives but was not further reduced in the normally menstruating females [15]. Notably, blood markers of muscle damage (myoglobin, creatine kinase, fatty-acid binding proteins) were all elevated by 48 hours of recovery for the males and females on oral contraceptives, but not the normally menstruating females, indicating a protective effect of estradiol on the secondary effects of damage [15]. The reduced secondary markers of muscle damage and enhanced recovery of neuromuscular function offer evidence that estradiol may attenuate muscle damage, yet the influence of damage on the protective effects of the RBE is unknown.

The purpose of this study was to investigate the relationship between menstrual cycle phase and neuromuscular impairments following an initial bout of eccentric exercise on the magnitude of the second bout. It was hypothesized that elevated levels of estradiol during the late follicular phase during the initial bout of eccentric exercise would attenuate deficits in neuromuscular function as compared with lower levels of estradiol in females on oral contraceptives, thereby resulting in a diminished repeated bout effect.

## 2. Material and Methods

### 2.1 Participants

Thirteen female participants (mass= 64.7 kg ± 13.1 kg, height = 70.8 cm ± 6.9 cm) ages 18-40 y were recruited from the University of Guelph community. Participants were required to be healthy and eumenorrheic. Participants were excluded from the study if they regularly participated in upper body exercise to reduce potential of a prior repeated bout effect. Additionally, participants were instructed to refrain from upper body exercise for the duration of the study. Procedures were all approved by the Human Research Ethics Board of the University of Guelph (REB: #23-08-003). Participants received verbal and written explanation of all aspects of the protocol. Written informed consent was obtained. Participants completed four study visits on separate days (Figure 1). Visits were divided into eccentric exercise protocol visits and 48hr follow-ups. The first eccentric protocol visit, and second eccentric protocol visit were separated by four weeks to allow for a recovery period. All baseline neuromuscular measures were completed on all visits and the eccentric exercise protocol was completed on visit 1 and 3. Participants were divided into two groups. One group (n=6) attended their first session during their late follicular phase of their menstrual cycle, corresponding to day 10-14 of their cycle estimated through cycle tracking with the onset of menses classified as day one. The second group (n=7) was tested on the placebo pills of second or third generation combined oral contraceptives.

**Figure 1.**
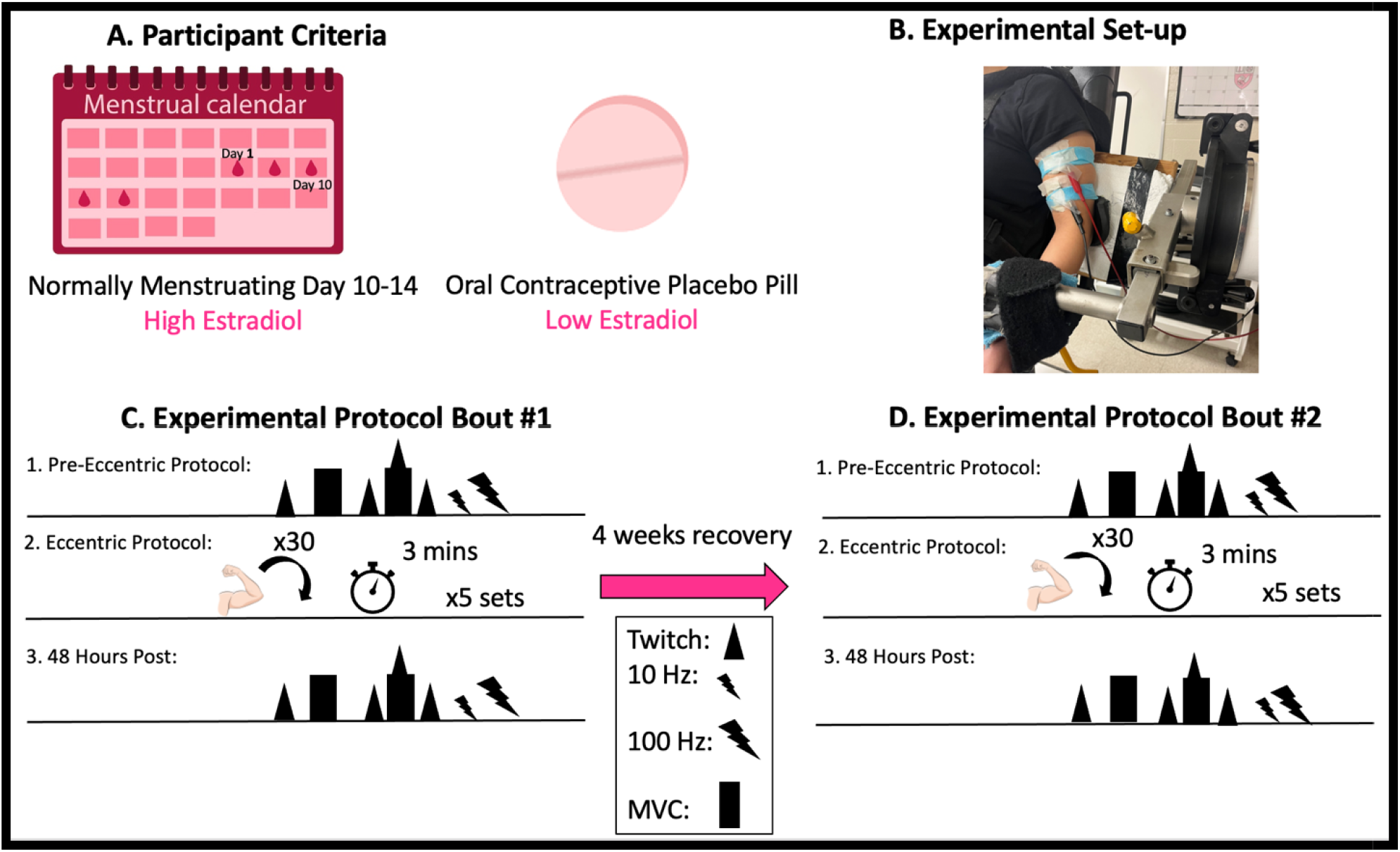
Experimental timeline. Baseline measures were conducted then participant performed 5 sets of 30 repetitions of maximal eccentric contractions at 180°/s. MVC, VA and PLFFD were then analyzed 48 hours after the eccentric protocol to examine effects of eccentric exercise induced neuromuscular impairments. MVC = maximal voluntary contraction; VA = voluntary activation; PLFFD = prolonged low frequency force depression

### 2.2 Experimental Set-Up

Participants were seated in a HUMAC NORM dynamometer (CSMi Medical Solutions, Stoughton, MA) and secured at the waist and shoulders with an adjustable 4-point non-elastic harness to limit whole body movement during each study visit. The left shoulder was secured further with a 5-inch wide Velcro strap running across the body (Figure 1). The elbow axis of rotation was placed in line with the axis of the dynamometer. The non-dominant left arm was securely attached in a supinated position with the arm at 110° of elbow flexion (terminal elbow extension being 180°) for all static contractions. The range of motion of the arm was 50° to 140° excursion [16]. Electrical stimulation was delivered with two custom stimulatory pads which were constructed by multiple layers of aluminum foil secured in a paper towel. The top of the pad was covered in tape while the bottom layer was soaked in water then covered with conductive gel. Alligator clips were secured to the aluminum foil so that the proximal pad acted as a cathode and the distal pad as an anode.

Optimal twitch torque was determined using percutaneous pad stimulation on the proximal and distal ends of the biceps brachii muscle belly using a high-voltage stimulator (DS7AH, Digitimer, Welwyn Garden City, Hertfordshire, UK). Torque (Nm) and stimulus data were all collected at 1000Hz using a 12-bit-analog-to-digital converter (PowerLab System 16/35; ADInstruments, Colorado Springs, CO, USA), and analyzed with Labchart (Version 8; Labchart, Pro Modules 2014, Colorado Springs, CO, USA) software. The current that elicited the maximal twitch force was found for each participant (pulse width of 1000 µs) by stimulating participants until the twitch force no longer increased with increasing current (DS7AH; Digitimer, UK). This stimulation current was increased by 20% and used during the interpolated twitch technique to estimate voluntary activation, described below. An isometric maximal voluntary contraction (MVC) was then performed. During each MVC, participants received strong verbal encouragement and visual feedback of torque production on a computer monitor positioned approximately 1 m from the participant. Torque guidelines were displayed to provide motivation for participants to achieve a higher maximum torque with each attempt. The participants were given a minimum of three attempts separated by three minutes of rest. The objective criteria used to deem an MVC as a maximal effort was 1) no further increase in torque between attempts and 2) voluntary activation of >90%. Voluntary activation (VA) was determined as follows: %VA = [1 - (Superimposed Twitch Torque/Potentiated Twitch Torque)] x100%. Participants were given two additional attempts if they felt they could obtain a higher torque value after the initial 3 attempts. The MVC torque was reported as the highest 500 ms average torque from a single attempt prior to the superimposed twitch.

In a similar manner to determining peak twitch torque, the current for tetanic stimulation was determined by increasing current during 1-s 100 Hz trains to evoke 25% of the participants MVC torque amplitude. After 3 min rest, this current was then used for 10 Hz and 100 Hz, in that order, to minimize potentiating effects of a higher stimulation Hz prior to a low stimulation Hz. Peak torque was determined as the highest torque reached during the contraction and was recorded for each frequency. The ratio of 10 Hz to 100 Hz (10:100 Hz) was classified as prolonged low frequency force depression (PLFFD).

Participants then completed five sets of 30 maximal isokinetic eccentric contractions at 180°/s of the elbow flexors separated by 3 minutes of rest between sets. Each eccentric contraction began at 50° flexion and went through a 90° range of motion. The participant was instructed to allow the dynamometer to passively return the elbow to 50° flexion at 30°/s. Strong verbal encouragement and visual feedback was provided through each contraction to encourage maximal effort. All neuromuscular assessments noted above were then performed 48 hours following the eccentric protocol. Self-reported soreness (visual analog scale of 1-10 cm) were measured at baseline and throughout recovery. “No pain” (0 cm) and “Severe pain” (10 cm) served as the anchors. The entire protocol was repeated in an identical manner 4 weeks later for Bout 2. The effect of bout is defined as the magnitude of the RBE.

### 2.3 Data Analysis

Statistical analyses were performed using IBM SPSS Statistics (v26). A one-way ANOVA was used on absolute data [Group (late follicular phase, placebo pills)] to access the impact of menstrual phase and placebo pill on baseline neuromuscular function prior to the eccentric protocols. A two-way repeated measures ANOVA [Group (late follicular phase, placebo pills) × Bout (Bout 1 48 h, Bout 2 48 h)] was used to assess the effects of menstrual phase and placebo pill on the magnitude of the repeated bout effect at 48 hours after Bout 1 and 2 to detect differences in peak twitch torque, MVC torque, VA, and PLFFD relative to baseline between the normally menstruating females and females on oral contraceptives. Additionally, to highlight changes after the eccentric protocol, a two-way repeated measures ANOVA [Group (late follicular phase, placebo pills) × Time (Bout 1 pre, Bout 1 48 h)] was used to compare the effects of group and time for Bout 1 & 2. Holm-Sidak post-hoc tests was utilized for all pairwise comparisons with an *α*□=□0.05. Data in text and figures are presented as mean ± SD.

## 3. Results

### 3.1 Baseline Group Differences

Prior to both eccentric exercise bouts, there were no significant differences between normally menstruating females and females on combined oral contraceptives (Figures 2 & 3) for: MVC torque (p=0.728), VA (p=0.444), soreness, electrically evoked torque at 10 Hz and 100 Hz, and the 10:100 Hz torque ratio (p>0.05).

**Figure 2.**
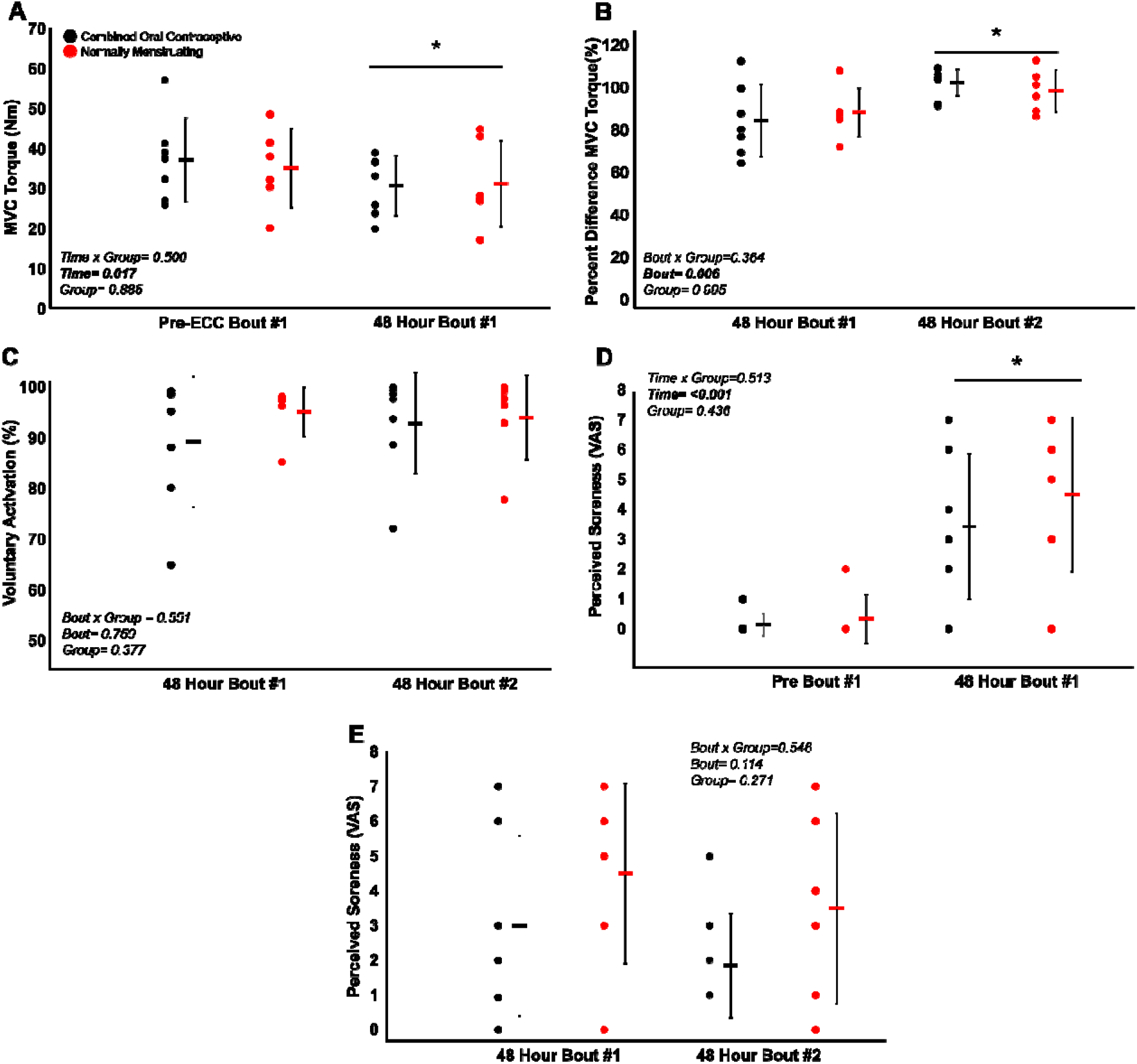
Neuromuscular function following bout 1 and 2. Changes in **(A)** Pre-eccentric exercise MVC, (**B)** MVC percent difference, **(C)** VA (%), **D)** soreness pre-eccentric exercise and 48 h following for Bout 1 and **(E)** soreness 48 h following the eccentric protocol for both bouts. Combined oral contraceptive females are in black and normally menstruating females are in red. Data are presented as mean ± SD. MVC = maximal voluntary contraction; VA = voluntary activation.

### 3.2 Eccentric Exercise Induced Neuromuscular Impairments and the RBE

Despite no difference in VA (p=0.750), there was a ∼15 % decrease in MVC torque for both groups (p<0.05) 48 hours following Bout 1 (Figure 2a) indicating significant long-lasting muscle weakness. Similarly, both groups had a significant increase in soreness (p<0.001) by 48 hours (Figure 2d). Most notably, there was a robust RBE following Bout 2, as indicated by no reduction in MVC following the eccentric exercise protocol (Figure 2b), indicating a complete protection against exercise-induced weakness. There was no effect of group (p=0.995), nor interaction (group × time; p=0.364) for MVC torque. Soreness across bouts was not significantly different (p=0.114). There was no difference in soreness between groups (p=0.271), or a Bout × group interaction (p=0.546; Figure 2e). Similar to MVC, there was a significant RBE for 100 Hz torque (p<0.05; Figure 3b), with no interaction (p=0.629) or effect of group (p=0.987). Meanwhile, 10 Hz torque displayed no significant Bout × group interaction, effect of Bout (p=0.477), nor effect of group (p=0.945). When comparing the ratio of 10:100 Hz, there was no effect of group (p=0.663), Bout (p=0.664), nor an interaction (Bout × group; p=0.593) indicating no significant PLFFD (Figure 3c).

**Figure 3.**
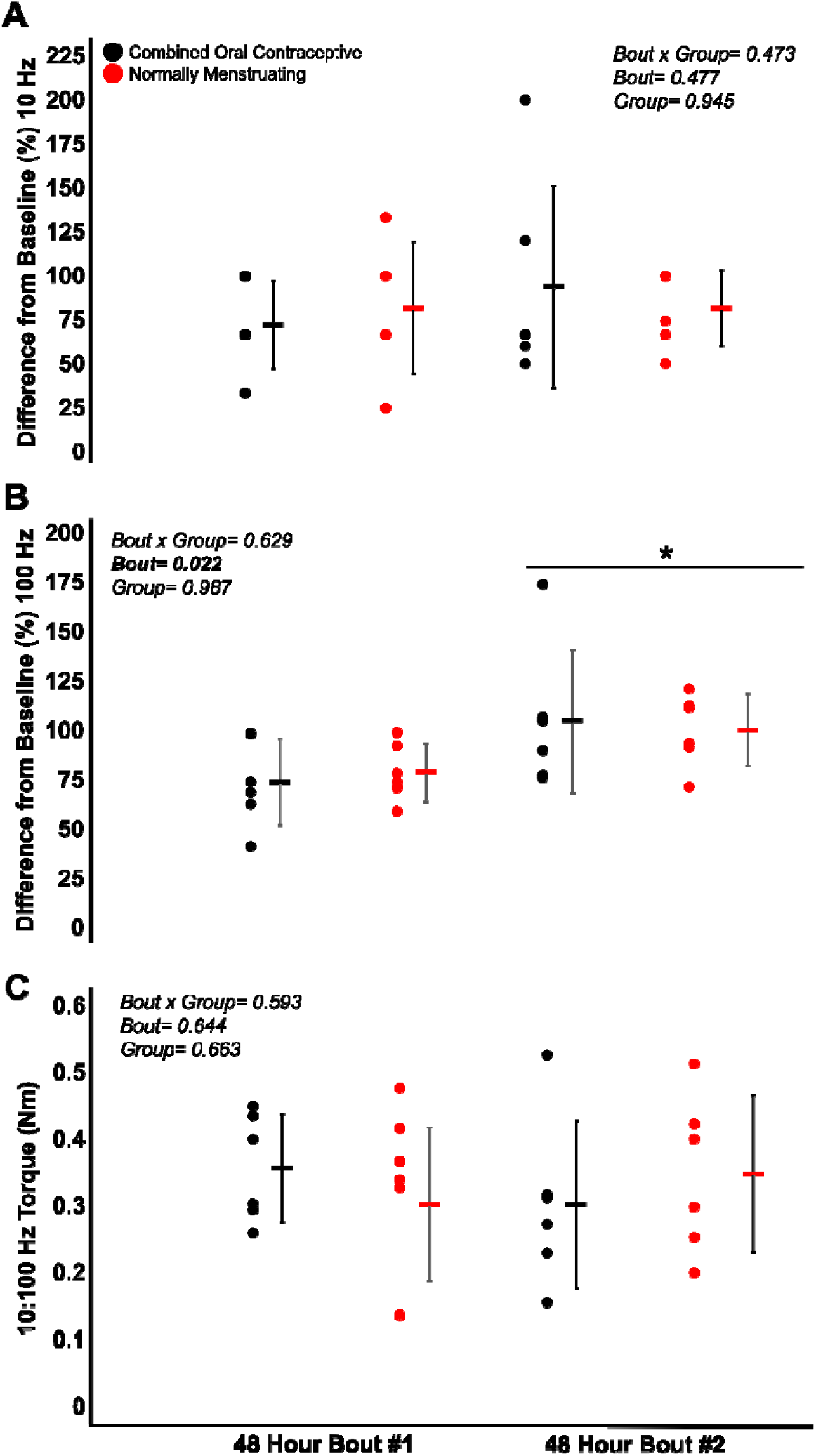
Prolonged low frequency force depression. Response at **(A)** 10 Hz, **(B)** 100 Hz, and **(C)** 10:100 Hz at 48 h following the eccentric protocol. Combined oral contraceptive females are in black (N=6; 1 participant did not complete the electrically evoked contractions) and normally menstruating females are in red. Data are presented as mean ± SD. PLFFD□=□prolonged low-frequency force depression.

## 4. Discussion

The present study investigated the effects of menstrual cycle phase and contraceptive use on eccentric exercise-induced neuromuscular impairments and the RBE. Contrary to our hypothesis, we did not observe any effects of menstrual cycle phase and oral contraceptive use on eccentric exercise-induced weakness and soreness. As well, there was a robust RBE which provided similar protection against eccentric exercise-induced muscle weakness regardless of assumed high/low estrogen levels throughout the menstrual cycle.

### 4.1 Eccentric Exercise-Induced Neuromuscular Impairments

The high-intensity eccentric exercise protocol induced long-lasting impairments in neuromuscular function (Figure 2). MVC torque was reduced for both groups 48 hours following Bout 1. Therefore, neuromuscular impairments were not attenuated to a greater extent in the late follicular phase of the menstrual cycle as hypothesized. This finding is consistent with Clarkson and Hubal (2021) who demonstrated that despite animal models showing a clear reduction in muscle damage in females, in humans differences in eccentric-exercise induced muscle damage between the sexes appears to be equivocal [17]. That said, females appear to have a more rapid onset of the inflammatory response which is mediated quicker than in males [17]. Neutrophils may be more rapidly alleviated in females, however, there appears to be a negligible effect of this attenuation on the extent of muscle damage in humans [17]. Additionally, there were no significant differences for biomarkers of muscle damage after eccentric exercise in females and males [18]. Overall, fluctuating sex hormones in healthy young females do not appear to mediate/alter eccentric exercise-induced muscle damage.

### 4.2 The Repeated Bout Effect

There was a significant RBE in both normally menstruating females in the late follicular phase with presumably high estradiol levels and females on combined oral contraceptives with low estradiol levels, such that the neuromuscular system was completely protected following eccentric-exercise Bout 2, with no decline in MVC (Figure 2b). The RBE is mediated by adaptations in the neuromuscular system following the initial bout of eccentric exercise which offers a protective effect against the subsequent bout. Various mechanisms are suggested to contribute to the magnitude of the RBE. These include neural adaptations, extra-cellular matrix remodeling, and the addition of serial sarcomeres thereby distributing the strain across more contractile units [19]. In our study, VA remained consistent and high throughout the duration of the study. Therefore, one’s ability to voluntarily activate the muscle near maximally was unlikely to be contributing to the RBE. Additionally, there was no difference in PLFFD between bouts indicating calcium release and myofibrillar calcium sensitivity was not different across bouts [20]. Therefore, owing to a similar VA across both bouts, and no difference in PLFFD, neural activation and changes in calcium release were unlikely to contribute to the magnitude of the RBE in our study. Instead, and beyond the scope of the present study, the main mechanism contributing to the RBE in our study was likely changes in sarcomere mechanics, extra-cellular matrix remodeling, and adaptive changes to the muscle-tendon unit [19] which requires further investigation.

### 4.3 Menstrual cycle phase and performance

There was no observed effect of menstrual cycle phase and oral contraceptive use on neuromuscular function following maximal eccentric exercise. Therefore, high levels of estradiol present in females in the late follicular phase and low levels in females on combined oral contraceptives do not impact the extent of muscle damage following unaccustomed eccentric exercise, resulting in no impact on the magnitude of the RBE. It is well understood that estradiol attenuates neutrophils and mitigates the extent of muscle damage in rodent studies such that female rats demonstrate reduced inflammation, β-glucuronidase activity and creatine kinase activity following muscle damage [21-23]. However, our understanding of how high levels of estradiol attenuate muscle damage in rodents as compared with humans is under-researched. Our current consensus on the impact of estradiol on neuromuscular impairments in humans is equivocal. Several studies suggest that specific menstrual cycle phases, and corresponding hormone levels may alter MVC torque while others observe no effect [24]. This discrepancy between rodents and humans is likely attributed to more extreme and quantifiable levels of estradiol in rodent models. Rodent studies often compare females to males [21-23]. However, males have differing concentrations of sex hormones compared to females. Other studies remove the rodent’s ovaries then inject exogenous estradiol [21-23]. Importantly, ovariectomies also result in a decline in progesterone [25], which is antagonistic to estradiol, and consequently progesterone’s removal heightens the action of estradiol [26]. Therefore, these models are not highly transferable to humans and likely explain some of the variability in response to estradiol levels in the literature when comparing investigations of animals and humans. Overall, our findings support the growing literature that menstrual cycle phase has minimal to no impact on overall neuromuscular performance [1,27].

### 4.4 Methodological Considerations

One limitation that should be considered is that females’ exact levels of estradiol were not measured. Menstrual cycle days were counted by participants, with day one classified as the onset of menses. Menstrual cycle phases are highly variable; therefore, calendar counting may be inaccurate [28]. However, this limitation is largely mitigated as females on combined oral contraceptives maintain consistently lower levels of estradiol compared to normally menstruating females [29]. A previous study observed that a difference of 56 pg/ml in estradiol was significant enough to have a 2-fold attenuation in neutrophils [10]. Therefore, any potential variability in participants’ exact menstrual cycle day on our outcome measures is minimized. Another factor to consider is that not all females have 28-day cycles, therefore, estradiol levels may not be identical on both Bout 1 and 2. To mitigate this, we ensured that females taking combined oral contraceptives were on the placebo pill, guaranteeing the estradiol levels in females on oral contraceptives was still lower than the normally menstruating females even during the second bout as described above.

## 5. Conclusion

The maximal eccentric exercise protocol effectively induced long-lasting neuromuscular impairments in both females on oral contraceptives during the placebo phase and normally menstruating females in the late follicular phase. Consequently, there was a significant reduction in neuromuscular impairments following the second bout of the eccentric protocol compared to the first, indicating a RBE. However, there were no differences in the magnitude of the RBE between females who were normally menstruating with higher levels of estradiol as compared with females taking oral contraceptives with lower estradiol levels. Thus, the late follicular menstrual cycle phase with high estradiol and contraceptive use during the placebo pill with low estradiol does not significantly impact neuromuscular function following high intensity repeated eccentric contractions. In line with current recommendations [27], female athletes should prioritize training around how they feel, not their menstrual cycle phase to optimize performance outcomes.

## Acknowledgements

We would like to thank all the participants, Avery Hinks and Dr. Brian Dalton for comments on a previous version of this paper, and Pardeep Khangura for technical assistance.

## Ethics statement

Participants gave written informed consent prior to testing. All procedures were approved by the Human Research Ethics Board of the University of Guelph (23-08-003) and, with the exception of registration in a database, conformed to the Declaration of Helsinki.

## Data availability

Data are available from the corresponding author upon request.

## Author contributions

A.R. and G.A.P. conceived and designed research; A.R. performed experiments; A.R analyzed data; A.R. and G.A.P. interpreted results of experiments; A.R. prepared figures; A.R. and G.A.P. drafted manuscript; A.R and G.A.P. edited and revised manuscript; A.R. and G.A.P. approved final version of manuscript.

